# *C21orf2* mutations found in ALS disrupt primary cilia function

**DOI:** 10.1101/2022.02.28.482239

**Authors:** Mathias De Decker, Pavol Zelina, Thomas G Moens, Kristel Eggermont, Matthieu Moisse, Jan H. Veldink, Ludo Van Den Bosch, R. Jeroen Pasterkamp, Philip Van Damme

**Author notes:** Corresponding author. **CO-CORRESPONDING AUTHORS:** Philip Van Damme,. Mathias De Decker.

## Abstract

Amyotrophic lateral sclerosis (ALS) is a devastating progressive neurodegenerative disease that affects 1 in 400 people. Almost 40 genes have been associated with ALS, currently explaining about 15% of the ALS risk. These genes tend to cluster in certain disease pathways such as protein quality control, RNA metabolism and axonal function. Despite these advances, adequate treatments for ALS patients are still missing. In this study, we investigate the role of a newly discovered ALS gene, *C21orf2*, in ALS pathology. We show that C21orf2 is localized to the basal body of the primary cilium and plays an important role in ciliogenesis *in vitro* and *in vivo*. Knock down of C21orf2 also lowers cilia frequency and length in human iPSC-derived spinal motor neurons (sMNs). Furthermore, we show that intraflagellar transport is impaired, causing primary cilia to fail in transducing extracellular signals essential in the sonic hedgehog pathway. ALS-associated mutations in *C21orf2* lead to loss of binding to centrosomal proteins, loss of proper localization at the basal body and hereby prevent C21orf2 from carrying out its normal function in cilia by loss-of-function. Finally, we confirm that sMNs derived from iPSCs from ALS patients with *C21orf2* mutations display similar cilia dysfunction and have disturbed sonic hedgehog signaling. Collectively, our data reveal impaired cilia homeostasis as a novel disease mechanism at play in ALS, opening new avenues for further research.

## INTRODUCTION

Amyotrophic lateral sclerosis is a complex neurodegenerative disorder that is characterized by the progressive death of motor neurons in the spinal cord, brainstem and motor cortex leading to progressive muscle weakness ^1,2^. While clinical onset is usually between 45 and 70 years of age, both younger and older onset have been described^3^. The site where the first symptoms arise is often focal, but the disease spreads to other body regions and ultimately leads to death from respiratory failure. The average survival time of ALS is only 2-5 years after symptom onset and treatment options are limited, with only two FDA approved drugs (Riluzole, Edaravone) providing marginal effects on disease progression^4,5^. There is a strong genetic contribution to ALS etiology. About 5-10% of patients can be classified as having familial (fALS) as they have a first or second-degree affected relative. The remaining patients are classified as having sporadic (sALS)^6^. Currently, mutations in more than 40 genes have been reported to explain a large proportion of fALS cases^7,8^. In addition, pathogenic gene mutations are found in 5-17% sALS patients^9,10^. ALS is a heterogeneous disease, from a genetic, pathophysiological and clinical point of view. While many of the genes associated with ALS cluster in RNA biology, proteostasis and axon architecture/transport, no comprehensive disease mechanism has been found^11^. Recent advances in the genetics of ALS have paved the way for new studies into the disease mechanisms of motor neuron degeneration.

One of the newly discovered ALS genes is chromosome 21 open reading frame 2 (*C21orf2* also known as *CFAP410*)^12^. A recent Genome Wide Assiociation Study (GWAS) has identified mutations in the gene C21orf2 as an ALS risk factor. Rare heterozygous loss-of-function mutations and missense mutations have been associated with ALS and over 75% of the associated variants are predicted to be detrimental, based on bioinformatic modeling approaches^13^. The biological function of C21orf2 remains poorly characterized, yet the highly conserved leucine-rich-repeat domain (LLR), the ubiquitous expression and the reported autosomal recessive mutations found in axial spondylometaphyseal dysplasia, Jeune syndrome and retinal dystrophy, underscores its functional importance^15–18^. Moreover, reduced expression of *C21orf2* has been reported in the brain of Down syndrome patients, indicating the potential role it may play in neurons^19^. Many functional roles have been attributed to *C21orf2*, but studies for these suggested cellular processes are scarce.

Based on immunoprecipitation studies, *C21orf2* was found to interact with NIMA-related kinase 1 (*NEK1*). Interestingly, rare variants in *NEK1* have also been observed in 3%-5% of ALS patients^14^. C21orf2 has been implicated in actin cytoskeleton organization and the C21orf2-NEK1 interaction has been suggested to play a role in DNA repair^20,21^. Furthermore, it has been shown that C21orf2 is a substrate of NEK1 and stabilizes the kinase in a SCF^FBXO3^-dependent manner^22^. The same study also demonstrated that mutant C21orf2 undergoes excessive phosphorylation by NEK1, which increases its stability, resulting in turn in stabilization of NEK1^22^. *C21orf2* has also been proposed to regulate primary cilia, and siRNA-based functional genomics screen demonstrated that *C21orf2* is a possible interactor of the ciliome and showed it localizes to primary cilia structures in the photoreceptor cells of human, pig and mouse retinas^23,24^. Thus far cilia defects have remained largely unexplored in the context of ALS.

The effect of ALS-associated mutations on C21orf2 function remains unknown. In this study, we investigated the functional consequences of ALS-associated variants in *C21orf2* in cell lines and iPSC-derived motor neurons. *C21orf2* ALS-associated mutations altered primary cilia morphology on motor neurons and disrupted their crucial function, uncovering cilia dysfunction as a novel disease mechanism in ALS.

## RESULTS

### *C21orf2* is essential for ciliogenesis

As a first step to understand the cellular function of C21orf2, we performed a gene ontology analysis on reported interactors of C21orf2 (Fig. 1a, Extended data table 1). The top hit resulting from this analysis was cilium assembly, a pathway largely unexplored in ALS. Other significant ontologies (p < 0.05) were largely related to either ciliogenesis or microtubule stability (Fig.1b). To confirm these *in silico* findings, we performed immunocytochemistry (ICC) on human neuroblastoma cells (SH-SY5Y) to examine the localization of C21orf2 (Fig. 1c,e). As described previously, we used serum-starvation as a method to induce primary cilium formation on SH-SY5Y cells which were labelled with Arl13B (Extended Data Figure 1a)^23^. C21orf2 colocalizes with γ-tubulin, a centrosomal marker, localized to the basal body region of primary cilia. We hypothesized that the heterozygous nonsense and missense mutations found in ALS could perturb C21orf2’s role in primary cilia assembly and maintenance. To test this hypothesis, we designed three siRNAs targeting *C21orf2* and quantified primary ciliary length and frequency. All three of the tested siRNAs could significantly reduce protein level, with C21orf2-II being the most potent. We continued to use C21orf2-II in all of our following experiments (Extended Data Fig. 1b,c). Compared to scrambled-control treated cells, *C21orf2*-depleted cells displayed significantly reduced ciliary frequency and length (Fig. 1d-f). This observation was confirmed using different cilia markers (Extended Data Fig. 1a). The findings show the pivotal role of *C21orf2* for primary cilia assembly and maintenance and suggests that *C21orf2* ALS variants could potentially disrupt cilium homeostasis.

**Figure 1.**
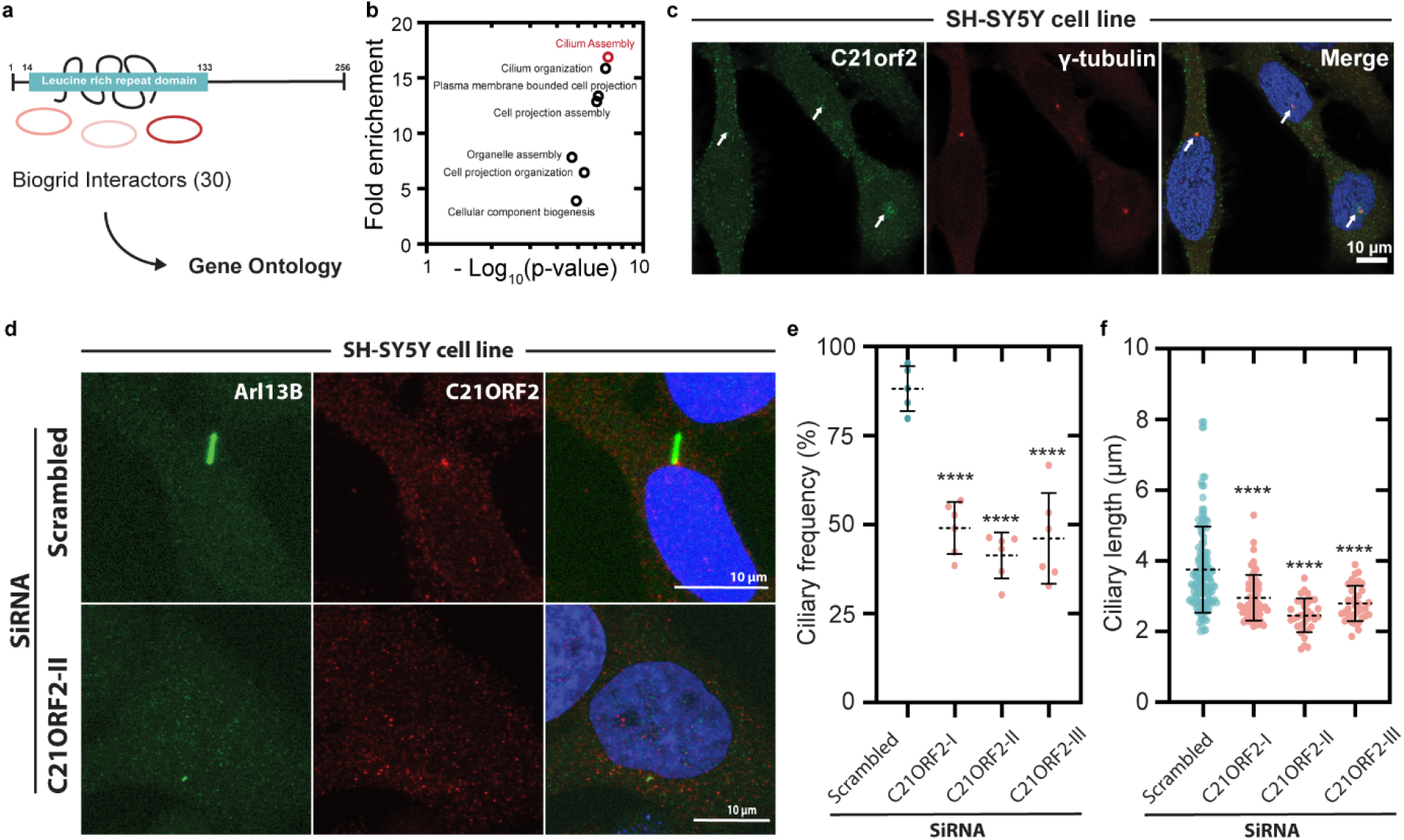
*C21orf2* regulates ciliary frequency and length. **a,** *In silico* study of the interactome of C21orf2 based on Biogrid interactors. **b,** Gene ontology prediction of the top hits of the interactome study. Red dot represent the best hit. **c,** C21orf2 and γ-tubulin staining of SH-SY5Y cells. White arrows show colocalization of C21orf2 with the centrosome marker (γ-tubulin). **d,** SH-SY5Y cells stained for Arl13B and C21orf2 after treatment with either scrambled (top row) or C21orf2 (bottom row) siRNA and starved in 0.5% FBS for 24h. **e,** Quantification of the ciliary frequency in SH-SY5Y cells after treatment with scrambled (blue) or C21orf2 (pink) siRNA. Mean ± s.d. (n = 5) are shown. Significance assessed by one-way ANOVA (F(3,19) = 32.02), followed by Dunnett’s post hoc test, **** represents p < 1.0 x 10^-4^. **f,** Quantification of the ciliary length in SH-SY5Y cells after treatment with scrambled (blue) or C21orf2 (pink) siRNA. Mean ± s.d. are shown. Individual data points (n >80) of three independent experiments. Significance assessed by one-way ANOVA (F(3,540) = 35.09) followed by Dunnett’s *post hoc* test, **** represents p < 1.0 x 10^-4^.

### DNA damage response is normal in C21orf2 depleted cells

Initial evidence linked *C21orf2* to the DNA damage repair mechanism^21^. This pathway is one of the proposed disease modulating mechanisms of ALS^25^. We therefore investigated if we could confirm these findings and potentially link *C21orf2* to this established ALS disease pathway. To determine if *C21orf2* is essential for DNA repair, we quantified the number of γH2AX positive foci in SH-SY5Y cells after treatment with a DNA damaging compound (0.1 mM H_2_O_2_). Cells with more than 10 γH2AX foci were counted as positive for DNA damage. 16 hours after treatment, no significant difference could be observed between C21orf2 depleted and control cells (Extended Data Fig. 1d,e). Additionally, we performed a comet assay in non-treated control and siRNA treated SH-SY5Y to detect double-strand DNA breaks (DSBs). In line with our previous findings, no significant increase in the tail moment could be observed in cell with lower levels of C21orf2 (Extended Data Fig. 1f,g). Based on our experiments, we did not find evidence that reduced levels of *C21orf2* modify the DNA damage response.

### Reduced C21orf2 levels disturb cilia functioning *in vivo*

*C21orf2* has homologues in nearly all of genome-sequenced vertebrates (Extended Data Fig. 2a). In addition, a *C21orf2* orthologue is also found in *Drosophila melanogaster* (*CG15208*). We used the GAL4-UAS system to reduce the levels of CG15208 *in vivo* via the use of an RNAi line expressed under the control of a ubiquitously expressed driver(tubulin-gal4) (Extended Data Fig 2b). After validating the knockdown levels using qPCR, we performed a phenotypic screen (Extended Data Fig. 2c). Flies with reduced levels of the *C21orf2* orthologue *CG15208* died shortly after eclosion because they immediately became stuck in the food. CG15208 knockdown flies showed no morphological changes, however a severely uncoordinated phenotype was observed. We additionally observed a large effect in the climbing ability in *CG15208* knockdown flies (Extended Data Fig. 2e). Both of these phenotypes are also observed in flies with defective mechanosensory transduction due to cilia dysfunction^26^. Mechanosensory neurons, located in type I sensory organs, are the only neurons in *Drosophila* equipped with specially modified cilia that transduce stimuli. Loss of cilia in the mechanosensory neurons results in morphologically normal but uncoordinated flies^27–29^. We used immunohistochemistry to visualize the mechanosensory neurons in the 2^nd^ antennal segment (also called Johnston’s organ). Ciliated structures were observed in normal control flies. In flies who had reduced levels of the *C21orf2 Drosophila* orthologue *CG15208*, these ciliated structures were no longer detectable (Extended Data Fig. 2g,h), while the overall neuronal distribution in the mechanosensory organ appeared normal. Additional evidence of defective cilia *in vivo* was found be examining a second ciliated structure. Most cells in the fruit fly have no cilia, we therefore focused on sperm cells and tested fertility^30^. Male flies with reduced levels of CG15208 were crossed with control females and progeny ratio was calculated based on the number of eggs in the tube. No progeny were observed when using CG15208 depleted males. Control flies on the other hand, had an egg to progeny ratio of almost 90% (Extended Data Fig. 2f). Based on our *in vitro* and *in vivo* findings we conclude that *C21orf2* is essential for primary cilia assembly and maintenance, and this stimulated us to explore if *C21orf2* variants found in ALS could potentially disrupt cilium homeostasis.

### *C21orf2* ALS variants cause ciliary defects

To assess the functional role of *C21orf2* in ALS, we generated constructs expressing *C21orf2* tagged with a C-terminal turbo-GFP (tGFP) and introduced missense mutations associated with ALS (V58L, Y68*, R106C, R172W, A255T)^12^. Mutations were chosen based on their allele frequency^31^ and predicted Combined Annotation Dependent Depletion (CADD) score^32^, a tool that estimates the deleteriousness of genetic variants (Extended Data Fig. 3a). Three of the variants chosen, were located in the highly conserved LRR domain of *C21orf2*. Substitutions of amino acids in these LRRs can change the stability of the protein or can disrupt the structural framework required for molecular interactions^33^. In an initial step, we investigated if the ALS-associated variants could act as gain of functions mutations. We therefore quantified ciliary length in cells expressing the different *C21orf2* constructs. Overexpression of wild type C21orf2 slightly increased ciliary length, whereas overexpression of mutant C21orf2 did not affect ciliary length compared to controls cells (Extended Data Fig. 3c). To ensure these differences were not explained by destabilization of the mutant protein we assessed protein expression using immunoblotting and observed no differences between the conditions (Extended Data Fig. 3b). Next, we wondered if ALS associated variants might lead to nonfunctional C21orf2 and therefore result in abnormal primary cilia in cells not expressing endogenous C21orf2. To test this hypothesis, we overexpressed the generated constructs in SH-SY5Y cells with reduced levels of C21orf2 and quantified ciliary frequency and length (Fig. 2a-c, Extended Data Fig. 3b). Whilst overexpression of wild type C21orf2 rescues C21orf2 knockdown (indicated on Fig. 2b,c as “no”), overexpression of LRR mutated constructs (indicated on Fig. 1b,c as “58”, “68”, “106”) could not rescue the reduced ciliary frequency nor the reduced cilia length. Furthermore, overexpression of variants (R172W and A255T) located in the conserved C-terminal domain and predicted to be deleterious mutations^13^, also could not rescue the reduced cilia frequency and length. These observations suggest that *C21orf2* ALS mutations impair an uncharacterized function in ciliogenesis. We hypothesized that these mutations may affect recruitment to the basal body. In support of this idea, we observed that mutated C21orf2 could not localize to the basal body of the primary cilia upon overexpression (Fig. 2a, Extended Data Fig. 3d). Quantifying the intensity of the GFP label over the intensity of the basal body marker, γ-tubulin, showed that all variants were more diffuse than wild-type *C21orf2* and failed to cluster at the basal body (Fig. 2d). Co-immunoprecipitations (CO-IP) with wild type and mutated constructs followed by western blot analysis showed that wild-type C21orf2 pulled down endogenous γ-tubulin but in mutated C21orf2, this interaction was abolished (Extended Data Fig. 3e). Furthermore in the same CO-IP, mutated C21orf2 failed to interact with centrosomal protein of 290kDA (CEP290), a known interactor of C21orf2 (Extended data table 1) and crucial protein in cilia development^34^. Collectively, these data indicate that the *C21orf2* mutations act as LOF mutations and are not able to rescue ciliary defects caused by the loss of C21orf2.

**Figure 2.**
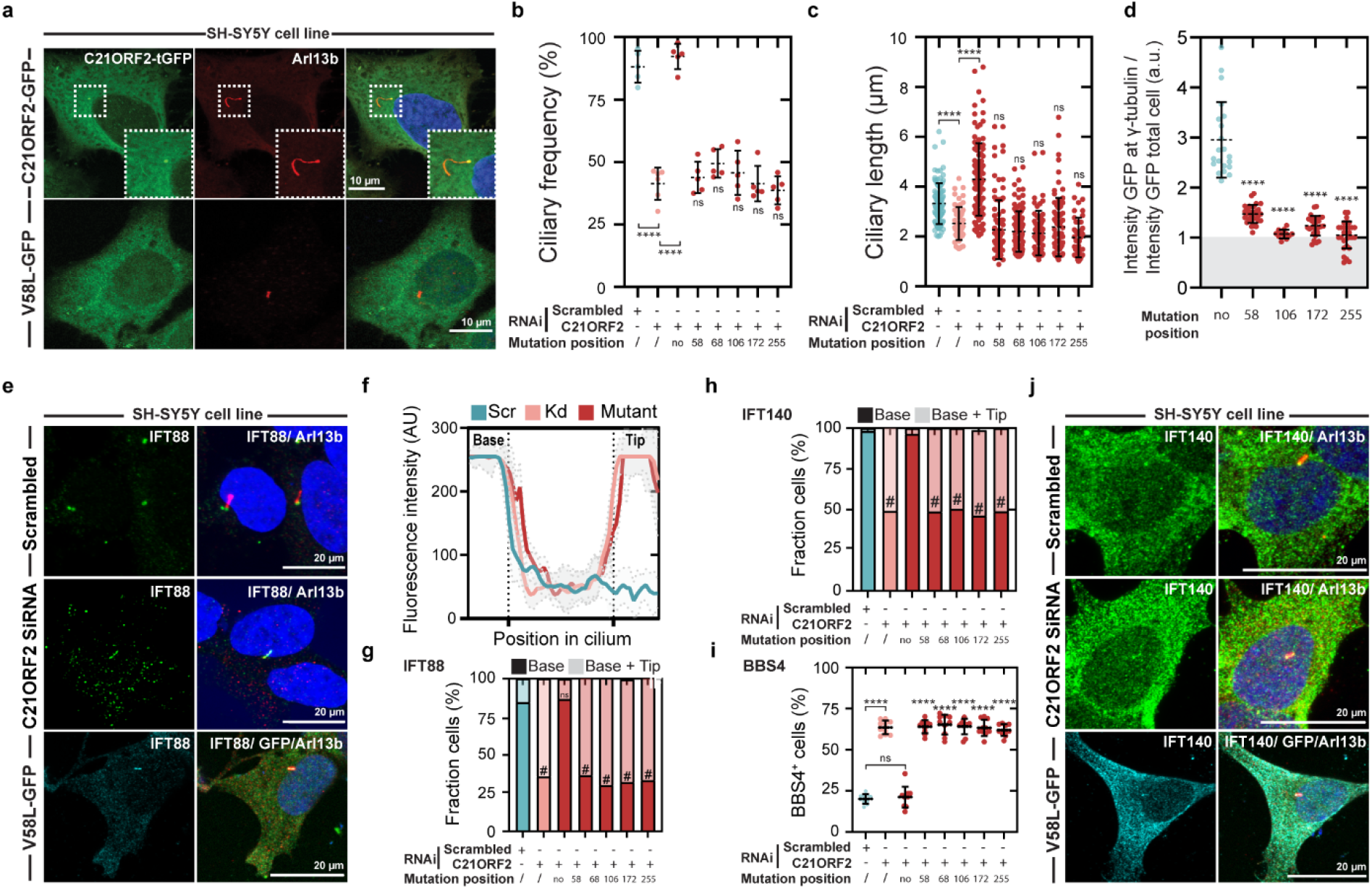
Mutations associated with amyotrophic lateral sclerosis are not able to rescue ciliary defects. **a,** SH-SY5Y cell transfected with *C21orf2-GFP* or *C21orf2-GFP* with a point mutation (V to L) on position 58. **b,** Quantification of the ciliary frequency (cells staining positive for the cilia Arl13b marker over the number of total cells) in the same setup as panel b. Mean ± s.d. (n=6) are shown. Significance assessed by one-way ANOVA (F(7,33) = 57,36) followed by Dunnett’s *post hoc* test, **** represents p < 1.0 x 10^-4^.**c,** Quantification of the ciliary length in SH-SY5Y cells transfected with the *C21orf2-GFP* wild type or *C21orf2-GFP* mutated at the indicated position. Mean ± s.d. are shown. Individual data points (n>90) from three independent experiments. Significance assessed by one-way ANOVA (F(7,664) = 53.47) followed by Dunnett’s *post hoc* test, **** represents p < 1.0 x 10^-4^.**d,** Quantification of the fluorescent intensity of GFP at the γ-tubulin^+^ location. Mean ± s.d. (n=20) are shown. Significance assessed by one-way ANOVA (F(4,117) = 46, 57) followed by Dunnett’s *post hoc* test, **** represents p < 1.0 x 10^-4^. **e,** SH-SY5Y cells, treated with scrambled or *C21orf2* targeting siRNA, stained for IFT88 and Arl13b (2 top rows). Bottom row, SH-SY5Y cells treated with *C21orf2* siRNA and transfected with *C21orf2-GFP* mutated at position 58. **f,** Fluorescence profile plot of IFT88 for the 3 conditions in panel e. Grey shade indicates sd’s of 4 independent experiments. **g,** Quantification of the fraction of SH-SY5Y cells showing IFT88 staining only in the base of the cilia (dark shade) or in the base and the tip of the cilia (light shade). Significance assessed by one-way ANOVA (F(6,75) = 209.7) followed by Dunnett’s *post hoc* test on the fraction of cells with a “base+tip” phenotype, # represents p < 1.0 x 10^-4^. **h,** Quantification of the fraction of SH-SY5Y cells showing IFT140 staining only in the base of the cilia (dark shade) or in the base and the tip of the cilia (light shade). Significance assessed by one-way ANOVA (F(6,42) = 159.0) followed by Dunn’s *post hoc* test on the fraction of cells with a “base+tip” phenotype, # represents p < 1.0 x 10^-4^. **i,** Quantification of fraction of SH-SY5Y cells staining positive for BBS4. Mean ± s.d. (n=10) are shown. Significance assessed by one-way ANOVA (F(7,72) = 180.8) followed by Dunnett’s *post hoc* test, **** represents p < 1.0 x 10^-4^. **j,** SH-SY5Y cells, treated with scrambled or *C21orf2* targeting siRNA, stained for IFT140 and Arl13b (2 top rows). Bottom row, SH-SY5Y cells treated with *C21orf2* siRNA and transfected with *C21orf2-GFP* mutated at position 58.

### C21orf2 is required for intraflagellar transport

A unique and highly regulated mechanism of primary cilia is intraflagellar transport (IFT). Proper biogenesis of cilia requires this specialized trafficking system for the movement of proteins into and out the organelle^35^. Moreover, defects in IFT have been linked to several human diseases^36^. To test if the observed morphological changes (shorter and less frequent cilia) causes disturbed IFT, we focused on two proteins; IFT88 and IFT140. IFT88 and IFT140 are subunits of larger protein complexes (IFT-B and IFT-A) that are shown to regulate anterograde and retrograde IFT within the cilium, respectively^37–39^. IFT88 normally exits from the cilium and localizes to the basal body once the primary cilium is formed (Fig. 2e). In cells with reduced levels of C21orf2, IFT88 accumulates primarily at the tip of the cilium and forms a “bulb” phenotype (Fig. 2e-g). Overexpressing of wild type *C21orf2* reverts this aberrant accumulation, unlike overexpressing the ALS mutant *C21orf2* constructs (Fig. 2e-g). We observed a similar effect when we looked at anterograde transport, using IFT140, and at an essential cargo protein; basal body (BBsome) subunit 4, BBS4^40,41^, which are normally rapidly exported from the cilium after assembly (Fig. 2h-j). As such, we conclude that C21orf2 is required for proper IFT functioning and that mutant C21orf2 is not able to rescue this defective anterograde and retrograde transport.

### Shh signaling is perturbed upon *C21orf2* knockdown

Primary cilia serve a critical role in canonical sonic hedgehog signaling (Shh). Most of the key pathway components localize within the organelle or have been otherwise associated with the ciliary function^42–44^. Shh signaling is involved in multiple developmental and several adult homeostatic processes^45–47^. Consequently, precise regulation is required via controlled intraflagellar transport (IFT) of positive (SMO) or negative (GPR161; SUFU) regulators inside or out of the cilium^38,39,43,44,48,49^. The observed impaired anterograde and retrograde transport in *C21orf2* depleted cells, led us to the hypothesis that Shh signaling could be impaired. When we stimulated Shh signaling by administering the smoothened agonist SAG (200 nM, 24h), we observed SMO accumulation in the cilium, whereas in *C21orf2* siRNA treated cells, SMO did not accumulate inside the cilium (Fig. 3a,b). Next, we investigated the negative regulator GPR161, which under basal conditions resides in the primary cilium and reverses its localization when Shh signaling is stimulated. In *C21orf2* knockdown cells treated with SAG (200 nM, 24h), GPR161 was not removed from the cilium (Fig. 3c,d). To further characterize the Shh signaling defects, we examined key transcription factors (*Gli1*, *Gli3*) downstream in the pathway. Gli3 is cleaved in the absence of Hh via a phosphorylation and proteasome dependent mechanism from Gli3 full length (Gli3FL) and Gli3 repressor (Gli3R)^50^. Hh stimulation represses this processing and results in a predominance of full length Gli3^51–53^. This effect is observed in control cells treated with SAG (200 nM, 24hr). In contrast, we could not replicate this effect in *C21orf2* knockdown cells (Fig 3e,f). Gli3R levels remained elevated after treated with SAG and Gli3FL levels were lower compared to control before and after treatment. Indicating an altered Gli3 processing and muted Hh signaling response. Next, we assessed the levels of *Gli1* after SAG treatment. *Gli1* is normally upregulated when the Shh signaling pathway is active, and we observed a robust induction of *Gli1* expression in control cells (Fig. 3f). However, in *C21orf2* knockdown SH-SY5Y cells, *Gli1* mRNA levels are not increased after SAG administration (Fig. 3f). To evaluate if more genes were misregulated in the Shh signaling pathway, we tested a gene panel of 84 candidates that are involved either in Shh signaling or in associated pathways (Extended data table 2). We treated control and *C21orf2* knockdown SH-SY5Y cells with 200 nM of SAG for 24h and extracted their RNA. Genes that were significantly (p<0.05) up or downregulated compared to control were summarized in fig.3g (Extended data table 2). Many of the key negative regulators of Shh signaling, SUFU; Gli3 and GSK3β, were significantly upregulated. In contrast, most downstream target genes were downregulated, suggesting a dormant Shh signaling pathway. Therefore, we can conclude that C21orf2 is required for responsiveness to the Shh pathway. Taken together, we showed that *C21orf2* mutations cause ciliary defects, which in turn, result in IFT malfunctioning and thereby, disrupt Shh signaling.

**Figure 3.**
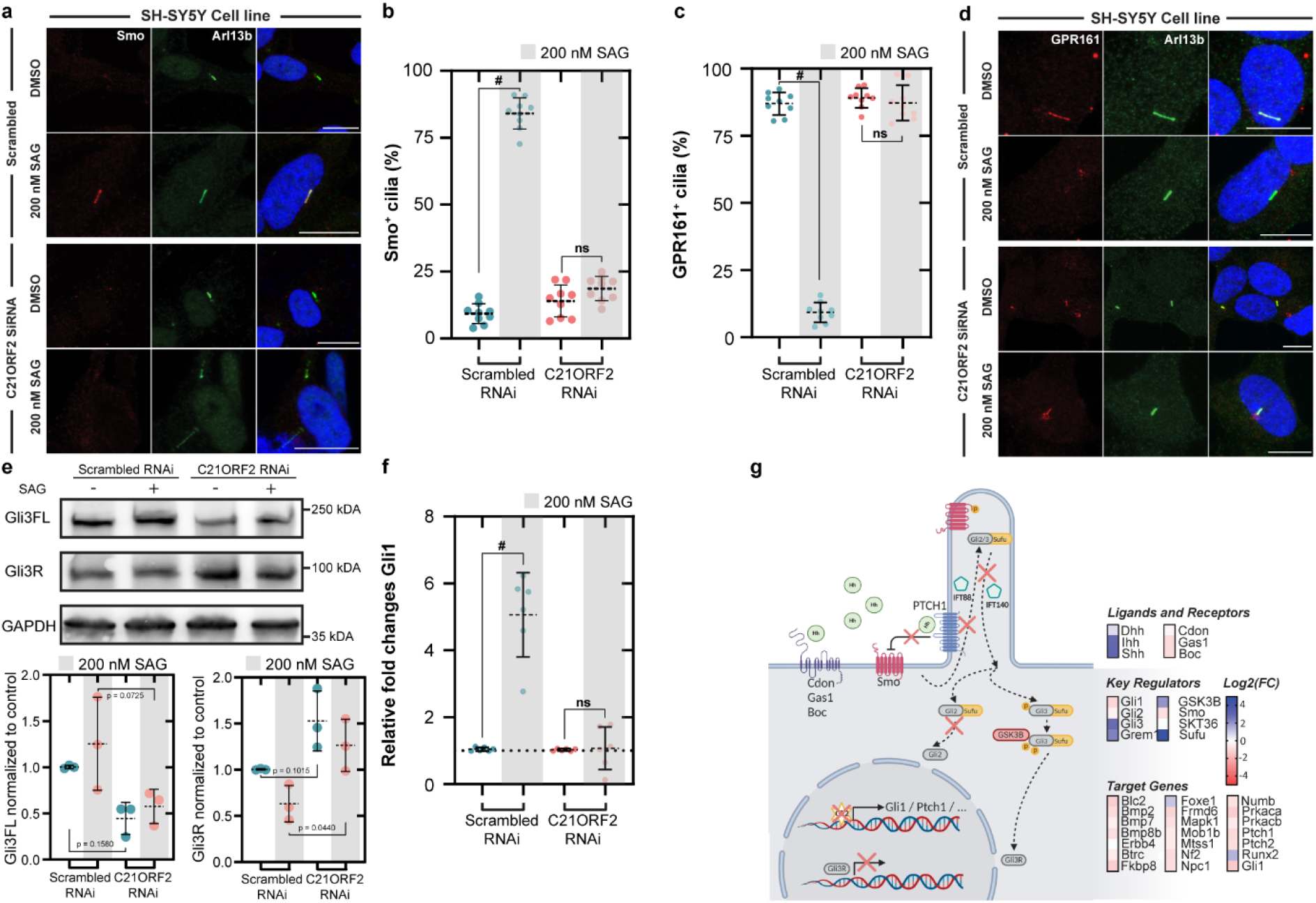
Loss of *C21orf2* results in defective sonic hedgehog signaling. **a,** SH-SY5Y cells with reduced levels of C21orf2 were treated with 200nM SAG for 24h and stained for Smo and Arl13b. Scale bar represents 10 μm. **b,** Quantification of the fraction of cells which stained positive for Smo. Significance assessed with unpaired two-tailed t-test, n = 10. Mean ± s.d. are shown. # represents p < 1.0 x 10^-4^. **c,** Quantification of fraction of cells which stained positive for GPR161 before and after treatment with 200 nM SAG for 24h. Mean ± s.d. are shown. Significance assessed with unpaired two-tailed t-test, n =9. Mean ± s.d. are shown. # represents p < 1.0 x 10^-4^. **d,** SH-SY5Y cells administered with siRNA that targets *C21orf2* or scrambled control, were treated with 200 nM SAG for 24h or vehicle control (DMSO) and stained for GPR161 and Arl13b. **e,** SH-SY5Y cells administered with *C21orf2* or scrambled siRNA, were treated with 200 nM SAG for 24h or vehicle control (DMSO). Gli3 full-length (Gli3FL) and Gli3 repressor (Gli3R) levels were examined using western blot analysis. C21orf2 depleted cells showed higher levels of Gli3R. Cells treated with *C21orf2* RNAi had lower levels of Gli3FL. Significance was tested by one-way ANOVA (F(3,8) = 5.288, Gli3FL, n =3) (F(3,8) = 7.850, Gli3R, n =3) followed by Šidák *post hoc* test. Mean ± s.d. are shown. **f,** qPCR for *Gli1* of SH-SY5Y cells treated with siRNA targeting *C21orf2,* in the presence of 200 nM SAG or DMSO. Significance assessed with unpaired two-tailed t-test, n = 6. Mean ± s.d. are shown. # represents p < 1.0 x 10^-4^. **g,** Summary of dysregulated genes associated with the sonic hedgehog pathway. Only significant genes are shown (cutoff p < 0.05). Significance tested using Geneglobe software (Qiagen). List of genes tested is compiled in extended data table 2.

### ISPC-derived motor neurons from ALS patients have defective cilia

We next studied if the primary cilia defects could be relevant in the context of ALS, as little is known about the role of cilia on motor neurons. To do so, we made use of human differentiated spinal motor neurons (sMNs) from fibroblast-derived iPSCs^54,55^. A first important observation was the presence of primary cilia on SMI-32 positive sMNs derived from control iPSCs (Fig. 4a). Although the cilia were less frequent than in immortalized cell lines, the ciliary length was roughly equivalent (Fig. 4c,d). Moreover, the measured ciliary frequency in our sMNs cultures were comparable with frequencies measured in primary motor neuron cell cultures from wild type mice^56,57^. To assess the effects of *C21orf2* ALS associated mutations, we employed a two-fold approach. First, we designed antisense oligonucleotides (ASOs) targeting *C21orf2*. Secondly, we generated sMNs from fibroblast-derived iPSC lines from two patients carrying an endogenous *C21orf2* mutation and two age-matched controls (Details of cell lines provided in Extended data table 3). To investigate the effect of reduced levels of C21orf2, we designed five ASOs targeting *C21orf2* and their matched scrambled controls. Their efficacy was tested using western blot and qPCR analysis and the most potent ASOs were used in follow-up experiments (Fig. 4b and Extended Data Fig. 4). The observed reduced levels of C21orf2 were comparable to the levels found when using siRNA in SH-SY5Y cells. We treated control sMNs with ASOs for ten days and quantified their ciliary frequency and length. sMNs treated with ASOs targeting *C21orf2* displayed significantly reduced ciliary frequency (Fig. 4c). In addition, knockdown of *C21orf2* caused a reduced ciliary length compared to MNs treated with scrambled controls (Fig. 4d). These results indicate the critical role of *C21orf2* in ciliogenesis, not only in SH-SY5Y cells as previously shown, but also in human motor neurons. To confirm this finding in a disease-relevant context, we differentiated motor neurons from iPSCs from *C21orf2* ALS patients carrying the V58L mutation (patient 1) and V58L/R60W compound heterozygous mutation (patient 2) in *C21orf2* (Extended data table 3). Again, we quantified ciliary length and frequency in sMNs that stained positively for the motor neuron marker SMI-32. Consistent with a loss-function in *C21orf2*, the sMNs from patients had significantly reduced ciliary frequency and length (Fig.4 e-g). To investigate if these cilia displayed functional defects, we treated our fully differentiated (38 days) patient and control motor neurons with 200 nM SAG for 24h and measured their Shh signaling response. We focused on three key targets in the Shh signaling pathway, which were misregulated in SH-SY5Y cells measured by the qPCR profiler array (Fig.3f). Control motor neurons treated with the Shh antagonist, SAG, responded by increasing *Gli1* mRNA levels. Motor neurons derived from ALS patients, on the other hand, failed to increase *Gli1* mRNA levels, suggesting an inactive Shh signaling pathway (Fig. 4h). Next, we examined two negative regulators of the Shh signaling pathway (*Gli3, SUFU*). Control sMNs showed reduced *Gli3* mRNA levels upon administration of SAG, but patient sMNs had an even slightly elevated *Gli3* transcript abundance. Similar observations were made when quantifying SUFU mRNA levels (Fig. 4h). In summary, sMNs derived from *C21orf2* ALS patients are unable to respond to extracellular Shh signaling stimuli due to defects in their primary cilium, suggesting that primary cilium defects are a novel disease mechanism in ALS.

**Figure 4.**
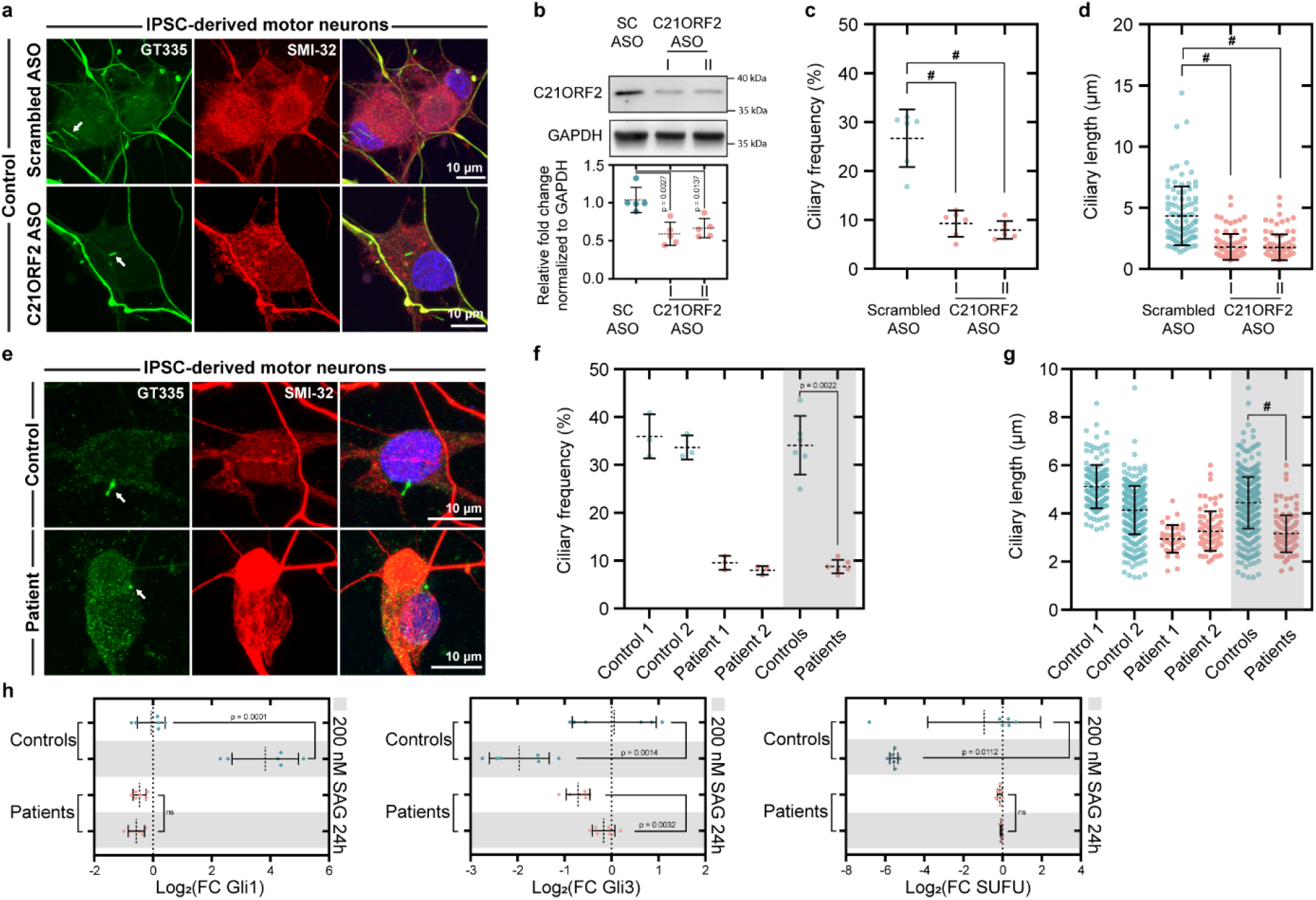
Motor neurons derived from IPSCs with reduced or mutated *C21orf2* have fewer and shorter cilia and cilia-associated signaling defects. **a,** Motor neurons derived from control iPSCs, treated with scrambled ASOs for 10 days (top row) or an ASO targeting *C21orf2*. Motor neurons were stained for SMI-32 (motor neuron marker) and GT335 (cilia marker). **b,** Western blot analysis of motor neurons treated with either scrambled or *C21orf2* ASOs. Mean ± s.d. are shown. Significance tested with one-way ANOVA (F(5,23) = 5.261, n = 5) followed by Dunnett’s *post hoc* test. **c,** Quantification of the ciliary frequency of motor neurons stained positive for SMI-32. Mean ± s.d. are shown. Significance assessed by one-way ANOVA (F(2,15) = 43.59, n = 6) followed by Dunnett’s *post hoc* test, # represents p < 1.0 x 10^-4^. **d,** Ciliary length measured in motor neurons that stained positive for SMI-32. Mean ± s.d. are shown. Individual data points (n > 90) of three independent experiments. Significance assessed by Kruskal-Wallis test followed by Dunn’s *post hoc* test. # represents p < 1.0 x 10^-4^. **e,** iPSC derived motor neurons from patients and aged-matched controls, stained for GT335 (cilia) and SMI-32 (motor neuron marker). **f,** Ciliary frequency measured in motor neurons which stained positive for SMI-32. Mean ± s.d. are shown (n = 6) and data was tested for significance using unpaired two-tailed t-test. **g,** Quantification of the ciliary length in motor neurons positive for SMI-32. Grey overlay shows grouped controls and patients. Mean ± s.d. are shown. Individual data points (n > 50) from three independent experiments. Significance tested using unpaired two-tailed t-test, # represents p < 1.0 x 10^-4^.**h,** qPCRs were used to assess transcript abundance of key regulators in the sonic hedgehog pathway. Motor neurons derived from iPSCs of controls and patients were treated with 200 nM SAG for 24h (grey overlay) or vehicle control. Mean ± s.d. are shown (n = 6) and data was tested for significance using unpaired two-tailed t-tests.

## DISCUSSION

Different possible disease mechanisms have been linked to motor neuron dysfunction and cell death in ALS, but none of these can fully explain the molecular biology of ALS. Recent large next-generation studies have resulted in a surge of the number of ALS-associated genes, which refine our understanding of ALS disease mechanisms. In this study, we investigate the role of new ALS-associated gene *C21orf2*. Our findings reveal a pivotal role for *C21orf2* in ciliogenesis and show the detrimental effects of mutated *C21orf2* on this crucial signaling organelle. Mutations associated with ALS in *C21orf2* acts as loss-of-function mutations by interfering with C21orf2’s interaction and localization with the basal body. Interestingly, motor neurons derived from ALS patients with *C21orf2* mutations display similar ciliary defects as observed upon knockdown of *C21orf2*. The functional consequences of defective cilia on motor neurons and potentially other cell types in ALS is still unknown and needs further investigation. An obvious downstream effect of perturbed cilia function is inhibition of the Shh signaling pathway, which is controlled by primary cilia. We showed that Shh signaling is abolished in motor neurons of ALS patients with *C21orf2* mutations, suggesting that this pathway is not only important in motor neuron differentiation, but also in motor neuron maintenance and function. A potential role of such an established developmental pathway in age-related neurodegeneration is emerging. Dysregulation of the Shh pathway has been reported in neurodegenerative diseases such as Alzheimer’s disease^47,58,59^, Parkinson’s disease^60,61^, Huntington’s disease^62^ and ALS^63–65^, demonstrating the importance of fully functioning Shh pathway, not only during development, but also during ageing. While further research is needed, we provide evidence that this dysfunction of this overlooked organelle is a contributing factor to the complex pathobiology in ALS. In conclusion, our results reveal a novel link between defective primary cilia and ALS pathology.

## SUPPLEMENTARY FIGURES

**Extended Data Figure 1.**
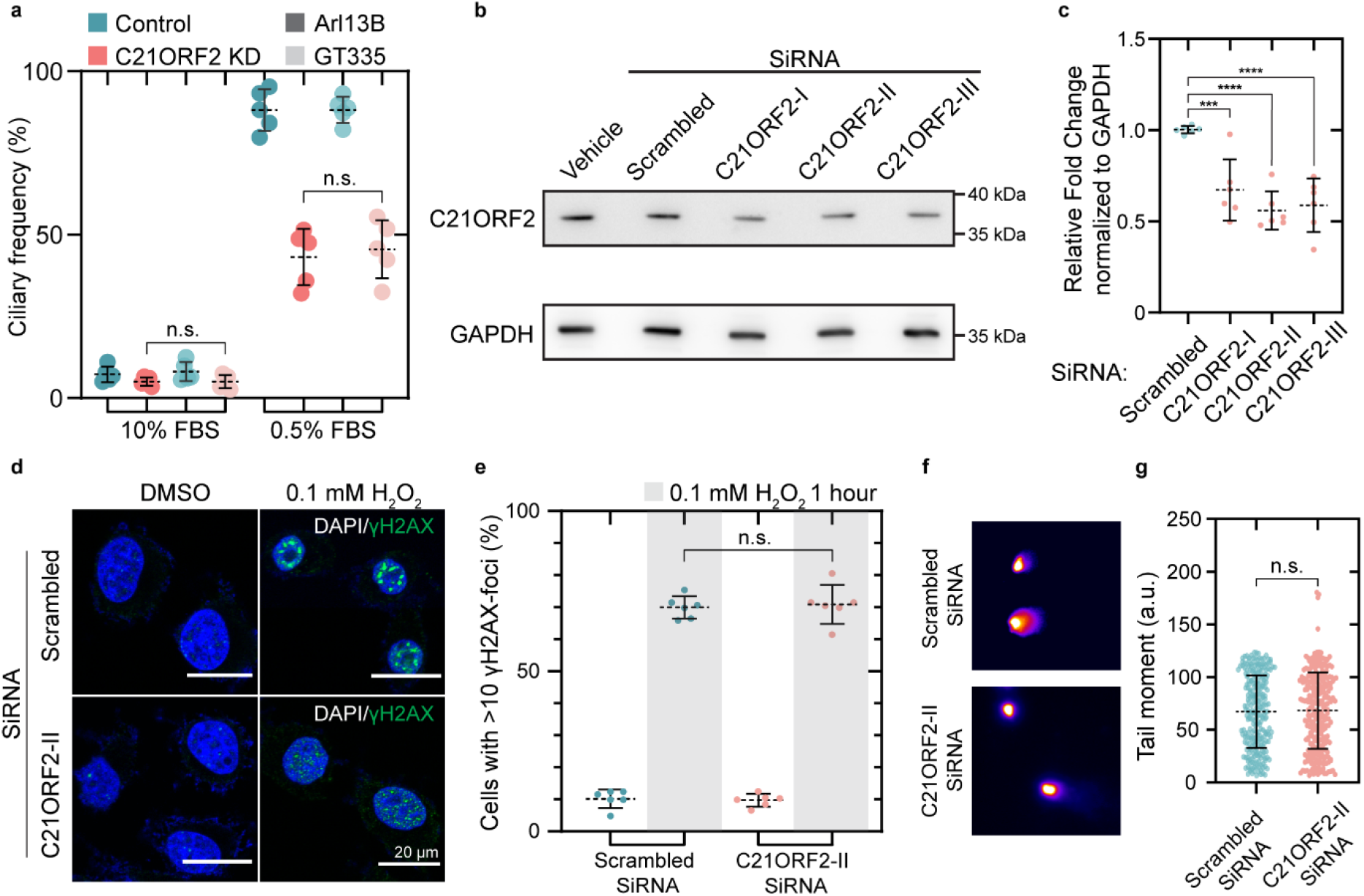
Quality control of cilia markers and siRNAs targeting *C21orf2.* DNA damage assays do not point towards an involvement of *C21orf2* in the DNA damage response. **a,** Scrambled or *C21orf2* siRNA treated SH-SY5Y cells were cultured for 36 hours in 10% or 0.5% FBS and ciliary frequency was quantified using ICC with 2 different cilia markers (GT335 or Arl13B). Significance was tested using unpaired two-tailed t-test (n = 5). P-value 10% FBS = 0.9937, P-value 0;5% FBS = 0.6733 **b,** Representative western blot of SH-SY5Y serum-deprived cells treated with siRNAs targeting *C21orf2*. **c,** Western blot analysis of C21orf2 levels normalized to GAPDH in SH-SY5Y cells treated with siRNA targeting *C21orf2* or scrambled control. Significance tested with one-way ANOVA (F = 16.31, n = 6) followed by Dunnett’s *post hoc* test. ***, **** represent p< 5.0 x 10^-4^ and p < 1.0 x 10^-4^ respectively. **d,** IHC of control and *C21orf2* depleted SH-SY5Y cells 16 hours after treatment with 0.1 mM H_2_O_2_, γH2AX (Green) labels sites of DNA damage. **e,** Quantification of percentage of SH-SY5Y cells with >10 γH2AX-foci, 16 hours after treatment with 0.1 mM H_2_O_2_. Significance assessed with unpaired two-tailed t-test (p = 0.7488 n = 6). **f,** Comet assay of SH-SY5Y cells treated with either scrambled SiRNA or *C21orf2* SiRNA. **g,** Quantification of the tail moment of the comet assay. Significance tested with unpaired two-tailed t-test (p = 0.7941). Individual data points (n > 250) from four independent experiments. Mean ± s.d. is shown.

**Extended Data Figure 2.**
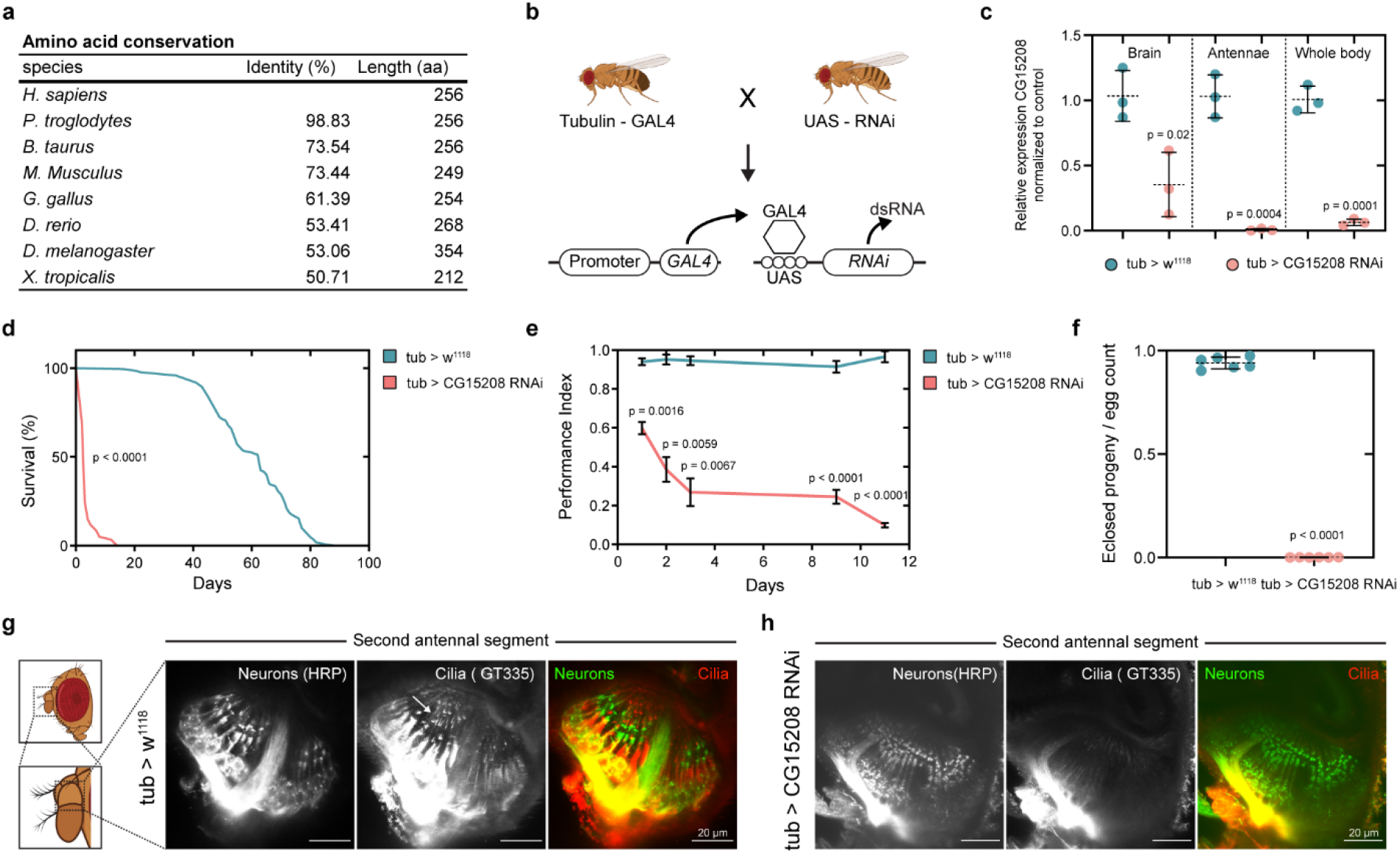
Knockdown of *C21orf2* fly orthologue results in cilia phenotype. **a,** Amino acid conservation among different species. **b,** Schematic overview of the GAL4-UAS system used to knockdown the *C21orf2* fly orthologue (*CG15208*). **c,** Expression levels of *CG15208* in different tissues before (control) and after knockdown). Significance tested by unpaired two-tailed t-test (n = 3). Mean ± s.d. are shown. **d,** Analysis of lifespan of control and flies with reduced levels of *CG15208.* Significance tested by Log-rank (Mantel-Cox) test. *(CG15208* knockdown, median lifespan = 3 days; control flies, median survival = 63 days) **e,** Negative geotaxis assay performed on *CG15208* knockdown and control flies. Significance tested via two-way ANOVA followed by Sidák’s multiple comparisons test. Mean ± SEM are shown. **f,** Assessment of male sterility in progeny of *CG15208* knockdown flies. Significance assessed with two-tailed t-test. Errors bars are mean ± s.d. **g,** Immunohistochemistry of mechanosensory neurons connecting to cilia in the 2^nd^ antennal segment (Johnston’s organ) in wild type pupae. Cilia indicated with arrow. **h,** Mechanosensory neurons lack cilia in *CG15208* knockdown flies. The distribution of neurons in the Johnston’s organ appeared normal but ciliated structures were no longer detectable.

**Extended Data Figure 3.**
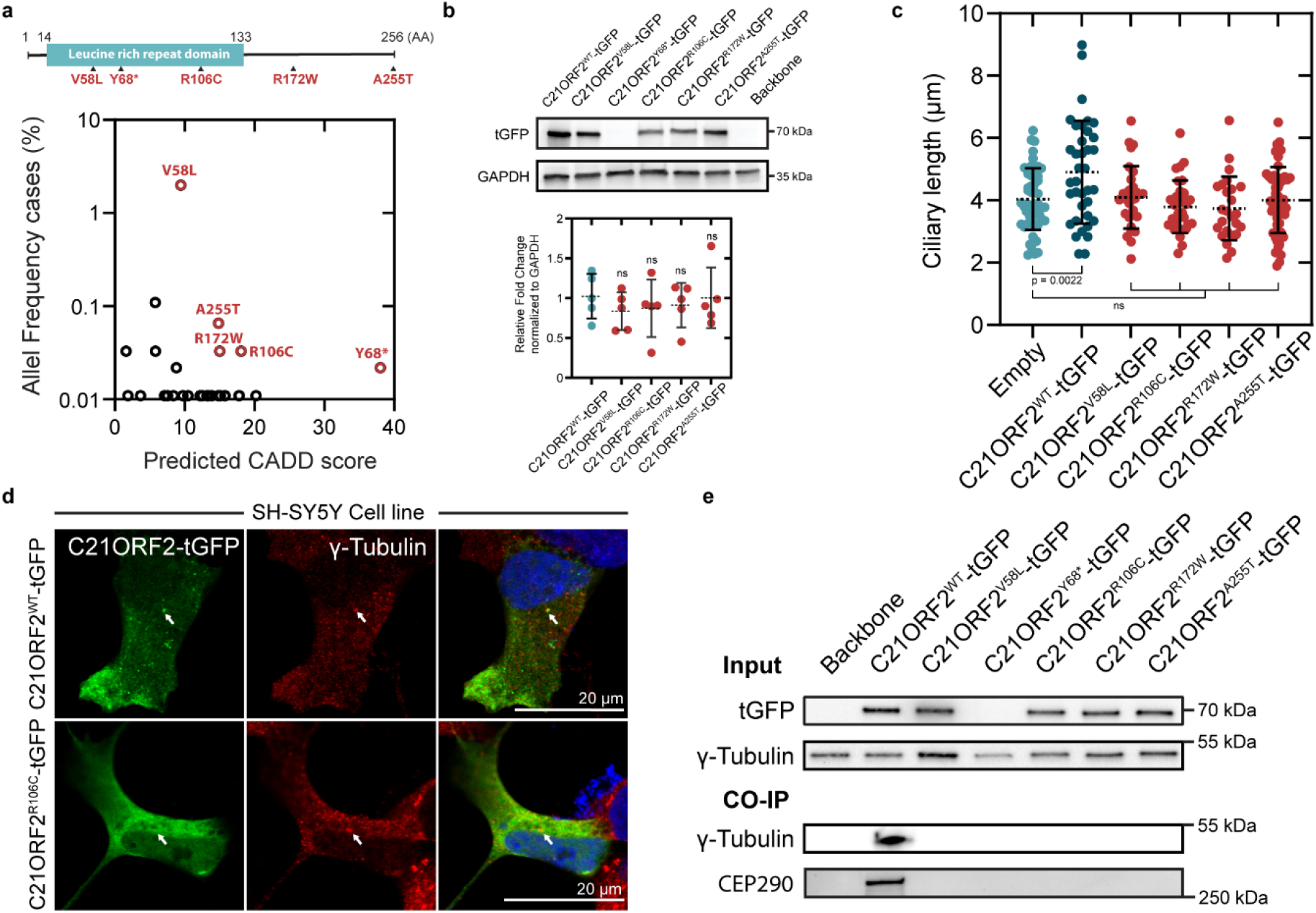
Characterisation of C21orf2 overexpression constructs. **a,** Scheme illustrating the locations of variants used to design overexpression constructs. *C21orf2* variants associated with ALS ranked based on allele frequency and predicted CADD score. **b,** Western blot analysis of lysates of SHSY-5Y cells transfected with C21orf2 constructs. Levels were assessed by the use of tGFP tag fused to the constructs. Quantification of the western blots. Levels are similar between wild type and mutant C21orf2 constructs. Significance tested by one-way ANOVA (F = 0.3451n = 5) followed by Dunnett’s *post hoc* test (ns, p > 0.05). Mean ± s.d. are shown. **c,** Quantification of ciliary length of SH-SY5Y cells transfected with *C21orf2* constructs. Overexpression of wildtype *C21orf2* increases ciliary length. Ciliary length in cells expressing mutated C21orf2 have similar lengths as controls (empty). Significance tested using one-way ANOVA (F(5,221) = 4.833) followed by Dunnett’s *post hoc* test. Individual data points (n > 30 of three independent experiments. Mean ± s.d. are shown. **d,** Mutated C21orf2 is not recruited to the basal body of primary cilia labeled with γ-tubulin. **e,** Co-immunoprecipitation experiment confirming the loss of interaction of γ-tubulin and CEP290 with ALS mutant C21orf2.

**Extended Data Figure 4.**
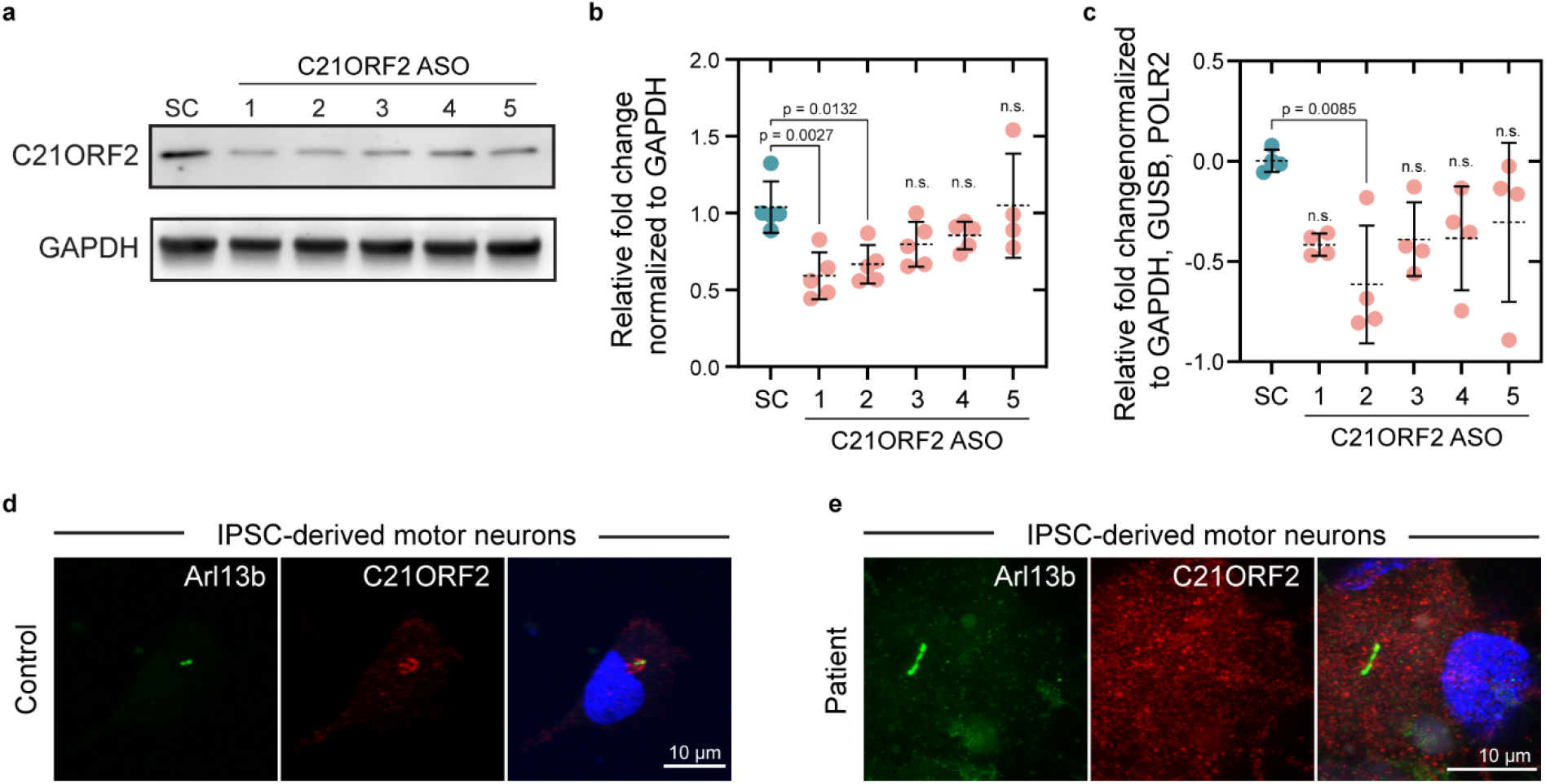
Evaluation of the efficacy of the designed ASO’s. **a,** Representative western blot of differentiated spinal motor neurons (sMNs) from control iPSCs and treated for 10 days with ASOs at a concentration of 50 nM. **b,** Quantification of western blot analysis. Significance was tested by one-way ANOVA (F(5,23) = 5.261, n = 5) followed by Dunnett’s *post hoc* test. **c,** qPCR analysis of sMNs treated for 10 days with ASOs at a concentration of 50 nM. Significance was tested by on-way ANOVA (F(5,18) = 2.784, n = 4) followed by Dunnett’s *post hoc* test. **d,** Representative image of C21orf2 location in control sMNs differentiated for 38 days. **e,** Representative image of C21orf2 location in patient (patient 2) sMNs differentiated for 38 days. Data are shown as mean ± s.d.

## ACKNOWLEDGEMENTS

We thank all members of the lab of Neurobiology for helpful discussions and suggestions, and Dr. Sandra Kling for help with the generation of iPSC lines. We would like to thank Dr Steven Boeynaems for feedback and advice. Illustrations were created with BioRender.com

## AUTHOR CONTRIBUTIONS

**M.D.D.**: Conceptualization, Investigation, Data Curation, Analysis, Writing Original Draft, Visualization. **P.V.**: Investigation. **T.G.M.**: Investigation, Supervision, Writing-Review & Editing. **K.E.**: Resources. **M.M.**: Investigation, Analysis. **J.H.V.**: Resources. **L.V.D.B.**: Supervision, Funding Acquisition, Writing-Review & Editing. **J.P.**: Supervision. **P.V.D.**: Conceptualization, Supervision, Funding Acquisition, Writing-Review & Editing.

## MATERIALS AND METHODS

### Cell culture and motor neuron differentiation

SH-SY5Y cells (Sigma, 94030304) were grown at 37°C in a humidified atmosphere with 5% CO_2_ in Dulbecco’s Modified Eagle’s Medium (Gibco, DMEM, F-12), high glucose, Glutamax + 10% Fetal Bovine Serum (FBS) and 2% pen/strep (Thermo Fisher Scientific). To induce ciliogenesis, cells were transferred to DMEM F-12 medium, Glutamax + 0.5% FBS and pen/strep for at least 16 hours. Cells were transiently transfected using Lipofectamine 3000 (Thermo Fisher Scientific) according to manufacturer’s instructions. Knockdown of *C21orf2* was achieved by administrating siRNA targeting *C21orf2* (Origene, SR300526) in a concentration of 50 nM using RNAiMAX (Thermo Fisher Scientific). Experiment set-up using siRNA or overexpression constructs followed the same timeline: cells were seeded on cover slips (#1.5) in medium with 10% FBS for 24 hours. siRNAs or plasmids were added to the medium and incubated for at least 48 hours or 24 hours, respectively. Afterwards, cells were transferred to medium supplemented with 0.5% FBS and starved for at least 16 hours. The control iPSC cell line used in ASO experiments was purchased from Sigma-Aldrich (iPSC Epithelial-1, IPSC0028). Two control and patient fibroblast lines were reprogrammed by Sendai virus-mediated expression of embryonic stem cell specific genes as well as addition of embryonic stem cell defining factors (Klf4, Oct3/4, Sox2, and cMyc). Absence of Sendai-virus after reprogramming was checked with quantitative real-time polymerase chain reaction (qPCR). Pluripotency markers were checked by qPCR together with immunohistochemistry analysis. In addition, the formation of three germ layers were also evaluated by using the embryonic body formation assay.

All experiments were approved by the ethics committee of the University Hospital Leuven. iPSCs were kept in E-8 Flex medium (Gibco) with pen/strep (Thermo Fisher Scientific, 5000 U/ml, concentration used 50U/ml). Colonies were passaged every 4 days with ReLeSR (Stemcell technologies) after rinsing with Dulbecco’s phosphate-buffered saline (DPBS, Sigma-Aldrich) and plated on Matrigel, hESC-qualified (Corning). Mycoplasma contamination was routinely tested by Mycoalert mycoplasma detection kit (Lonza, LT07-318). iPSCs were differentiated into sMNs as previously described^54,66^. Mature sMNs (day 38) were used in all described experiments.

### Immunocytochemistry

Cells grown on coverslips (1.5#) were incubated for 20 minutes at room temperate (RT) in PBS with 4% paraformaldehyde (Sigma-Aldrich) and washed three time for 10 minutes with PBS. Cells were incubated for 1 hour at RT in PBS containing 0.1% Triton X-100 (Sigma-Aldrich, PBS-T) and 10% normal donkey (Sigma-Aldrich) or goat (Dako) serum. Primary antibodies were diluted in PBS-T and incubated overnight at 4°C. Cells were washed three times for 10 minutes with PBS-T before adding secondary antibodies diluted in PBS-T for 1 hour at RT (Extended Data Table 4). After three wash steps of 10 minutes in PBS, cells were incubated with Hoechst 33342 diluted in PBS (NucBlue Live ReadyProbes Reagent, Invitrogen). Coverslips were rinsed in PBS and mounted with Prolong Gold antifade reagent (Invitrogen). Confocal images were obtained with Leica SP8 DMI8 confocal microscope and images were analyzed, formatted and quantified using Fiji^67^. Measurement of cilia length was performed as previously described^68^.

### Western blot analysis

Cells were harvested on ice and flash frozen cell samples were maintained at −80 °C until further processing. Samples were lysed in RIPA buffer (containing 50 mM Tris, 150 mM NaCl, 1% (vol/vol) NP40, 0.5% sodium deoxycholate (wt/vol), 0.1% SDS (wt/vol) with protease inhibitors (Complete mini EDTA free, Roche Diagnostics, pH 7.6). Protein concentrations were determined using the microBCA kit (Thermo Fisher Scientific Inc.) according to the manufacturer’s instructions. For immunoblot analysis, samples (30μg) were resolved on 4 to 20% mini-protean TGX stain-free gels (Bio-Rad) and transferred to nitrocellulose membrane (Bio-Rad). Membrane was blocked with 5% nonfat milk (Bio-Rad) in TBS-T for 2 hours at room temperature and incubated with primary antibodies overnight at 4°C (Extended Data Table 4).

The next day, the membranes were washed three times with TBS-T and incubated for 1 hour with HRP-conjugated secondary antibodies (Dako). Proteins were detected by enhanced chemiluminescence reagents (Thermo Fisher Scientific) and detected with the ImageQuant LAS 4000 Biomolecular Imager (GE Healthcare Life Sciences).

### Co-immunoprecipation

Samples were collected on ice and lysed in ice-cold RIPA buffer (50 mM Tris, 150 mM NaCl, 1% (vol/vol) NP40, 0.5% sodium deoxycholate (wt/vol), 0.1% SDS (wt/vol) with protease inhibitors (Complete, Roche Diagnostics, pH 7.6). Cell debris was removed via centrifugation (20 min, >12000 RPM, 4°C) and supernatants was transferred to a new tube. Protein concentrations were determined and an equal amount of protein was mixed with TurboGFP-Trap Magnetic Agarose (Chromotek) for 1 hour at RT with agitation. Samples were processed according to manufacturer’s instructions and eluted by boiling the incubated beads in 2X SDS-sample buffer.

### Comet assay

Samples were collected using a cell scraper in ice cold 1X PBS (Sigma-Aldrich). Cells were transferred to a centrifuge tube and cell count was performed. After washing the cells in ice cold 1X PBS, cells were diluted to 1 x 10^5^ in ice cold PBS (Ca^2+^ and Mg^2+^ free). Samples were processed with the Comet Assay kit (Trevigen) according to manufacturer’s instructions. Samples were incubated in alkaline unwinding solution (200 mM NaOH, 1 mM EDTA, pH>13) and run in alkaline electrophoresis solution (200 mM NaOH, 1mM EDTA, pH > 13). Comet tails were scored using ImageJ based on intensity of the tail DNA content according to manufactures instructions (Trevigen, Comet assay Kit).

### Quantitative RT-PCR and pathway-focused gene expression analysis

RNA was isolated from snap frozen samples using the RNeasy kit (Qiagen). Reverse transcription was performed using Superscript III First-Strand Synthesis Supermix for qRT-PCR (Invitrogen). qRT-PCR was performed using probe-labeled primers (Thermo Fisher) on the Quantstudio 3 (Applied Biosystems). All samples were run in triplicate and quantified using qBase+ (Biogazelle). Housekeeping gene stability was checked using qBase+. Five housekeeping genes were screened for every sample type (immortal cell line, sMNs) and the top three were used to normalize expression levels. A list of primers can be found in extended data table 5. The Sonic Hedgehog pathway gene expression profile was assessed via RT^2^ profiler PCR array (Qiagen) according to manufacturer’s instructions. In short, RNA was collected from cell pellets with TRizol Reagent (Invitrogen) and quality of RNA was confirmed using Bioanalyzer (Aligent).Next, cDNA synthesis was performed using the RT^2^ First Strand kit (Qiagen). Finally, samples were mixed with RT2 SYBR Green Mastermix (Qiagen) and loaded on RT^2^ PCR profiler plates (Qiagen, PAHS-078Z). Results were analyzed using the GeneGlobe Data Analysis Center (Qiagen).

### Treatment of sMNs with ASOs

Locked Nucleic Acid (LNA) oligonucleotides for *C21orf2* were purchased from Qiagen. A complete list of the used ASOs can be found in extended data table 6. Spinal motor neuron progenitors were plated at low seeding density of 0.5 x 10^5^ cells per well (12 well plate) at day 10 of differentiation. At day 25, oligonucleotides were dissolved in DNase and RNase free water (Qiagen) and added to cultures. Normal maturation medium changes were performed every two days supplemented with 50 nM ASO concentration. Motor neurons were lysed or stained at day 38.

### Plasmids

*C21orf2* plasmid was purchased from Origene (Cat#: RG210047). Mutagenesis was performed using Quickchange II kit from Aligent with primers found in extended data table 5. Mutations were regularly validated using Sanger sequencing (LGC Genomics). Overview of all plasmids are summarized in extended data table 7.

### Fly culture conditions, stocks and transgenic lines

*Drosophila melanogaster* strains were maintained on standard yeast, cornmeal, and agar-based medium (5% glucose, 5% yeast extract, 3.5% wheat flour, 0.8% agar) in a 12 hr light/dark rhythm. The w1118 (Canton-S10) line was used as control. Crosses for life span, negative geotaxis and fertility assays were performed at 25°C. Progeny was collected daily and the flies were transferred to 25°C. GAL4 driver line yw*; P{tubP-GAL4}LL7/TM3, Sb1 Ser1 (5138) was obtained from the Bloomington Stock center. For all our experiments, the w^1118^ line crossed to the drivers was used as a control. The RNAi line for CG15208 was obtained from Vienna Drosophila Resource Center (VDRC ID 106724). The line was checked for 40D insertion using PCR and qPCR of Tiptop gene. No insertion or overexpression could be observed^5,6^.

### Quantitative RT-PCR of *Drosophila* samples

Flies were collected on day 2 and flash frozen in liquid nitrogen. For the whole body qPCR, 20 flies were homogenized in 1 ml TRIzol Reagent (Invitrogen) with a disposable plastic pestle. For the brain and antennae qPCR, tissues were dissected from 30 flies and homogenized using a disposable plastic pestle in 0.5 ml TRIzol Reagent. Samples were centrifuged at 12,000 x g for 10 minutes at 4°C. The fatty layer was removed and the cleared supernatant was transferred to a new tube. Chloroform was added in 1/5 ratio based on the TRIzol volume. After centrifugation (12,000 x g, 15 min, 4°C), aqueous phase was transferred to a new tube and precipitated with 0.5 ml 100% isopropanol. After centrifugation, RNA pellet was washed with 1 ml 75% Ethanol and resuspended in RNase-free water. Reverse transcription was performed using Superscript III First-Strand Synthesis Supermix for qRT-PCR (Invitrogen). qRT-PCR was performed using probe-labeled primers (Thermo Fisher) or using SYBR green on the Quantstudio 3 (Applied Biosystems). All samples were run in triplicate and quantified using qBase+ (Biogazelle).

### Lifespan Analysis

Newly eclosed flies were transferred into vials of standard food at a density of 10 flies per vial and were transferred to a new tube every 2 days. After each transfer, surviving flies were counted. The density per vial was controlled for, and female and male flies were separated. At least 100 flies per genotype were assessed.

### Negative geotaxis Assays

Female flies were placed in a transparent tube for 1 hour to acclimate at a density of 10 flies per vial. Flies were then tapped to the bottom of the tube and scored according to their ability to climb. Flies were counted as successful if they reached the 8 cm line in 10 seconds. The test was conducted twice per time point with an interval of roughly 15 minutes. 50-75 flies were tested. The performance index was calculated at the ratio of flies that were successful in the crossing the 8 cm line in 10 seconds, over the total number of flies in the vial.

### Sterility assay

Males with reduced levels of CG15208 were separated upon eclosion for 2 days. Isolated males were crossed with control 3-5 day old virgin females for 1 day in a ratio of 1 male to 2 virgin females. Afterwards the females were transferred to a new tube of normal food and were allowed to lay eggs for 1 day. Vials where either males or females died during mating or egg laying were excluded. After removing the females from the tube, the eggs were counted and allowed to develop for 10 days. Progeny rates were calculated by dividing the hatched flies to the number of eggs on day 0.

### Immunohistochemistry of the 2^nd^ antennal segment (Johnston’s organ)

Pupae (pupal stage P10-P13) were picked and heads were dissected and fixed in PBS 1X (Sigma) with 4% PFA (Sigma) for 30 minutes at room temperature. Heads were washed three times 5 minutes in PBS. Heads were transferred on a glass slide and antennae were dissected and stored in 0.65 ml low-binding tubes containing PBS 1X. Antennae were permeabilized with PBS 1X with 0.4% Triton X-100 (Sigma) for 1-2h at room temperature. Afterwards, antennae were incubated with primary antibodies diluted in blocking buffer for 48 hours. Samples were washed four times for 15 minutes with PBS and incubated with secondary antibodies for 48 hours at 4°C. After four washing steps of 15 minutes in PBS, antennae were mounted in prolong gold antifade mounting media (Invitrogen).

### Statistics

All data was analyzed using GraphPad Prism 9.2.0 and Excel. Statistical tests, p values, number of samples, replicates, and experiments are indicated in the figure legends.

